# Local Gene Duplications Drive Extensive NLR Copy Number Variation Across Multiple Genotypes of *Theobroma cacao*

**DOI:** 10.1101/2024.09.01.610724

**Authors:** Noah P. Winters, Eric K. Wafula, Prakash R. Timilsena, Paula E. Ralph, Siela N. Maximova, Claude W. de Pamphilis, Mark J. Guiltinan, James H. Marden

**Affiliations:** IGDP Ecology, The Pennsylvania State University, University Park, PA; Huck Institutes of the Life Sciences, The Pennsylvania State University, University Park, PA; Applied Science and Technology, Battelle Memorial Institute, Columbus, Ohio, USA; Department of Biology, The Pennsylvania State University, University Park, PA; IGDP Plant Biology, The Pennsylvania State University, University Park, PA; Department of Plant Science, The Pennsylvania State University, University Park, PA

**Author notes:** Corresponding author address: 505 King Avenue Columbus, Ohio 43201.

## Abstract

Nucleotide-binding leucine rich repeat receptors (NLR) are an essential component of plant immunity. NLR evolution is complex and dynamic, with rapid expansions, contractions, and polymorphism. Hundreds of high-quality plant genomes generated over the last two decades provide substantial insight into the evolutionary dynamics of NLR genes. Despite steadily decreasing sequencing costs, the difficulty of sequencing, assembling, and annotating high-quality genomes has resulted in comparatively little genome-wide information on intraspecies NLR diversity. In this study, we investigated the evolution of NLR genes across 11 high-quality genomes of the chocolate tree, *Theobroma cacao* L. We found 3-fold variation in NLR copy number across genotypes, a pattern driven primarily by expansion of NLR clusters via tandem and proximal duplication. Our results indicate local duplications can radically reshape gene families over short evolutionary time scales, creating extensive intraspecific variation and a source of NLR diversity that could be utilized to enrich our understanding of both plant-pathogen interactions and resistance breeding.

## Introduction

Plant immunity is principally composed of two layers. The first is formed by extracellular receptors that sense and respond to conserved molecular patterns, called pattern recognition receptors (PRR) (Gómez-Gómez and Boller 2000; Chinchilla et al. 2007; Zipfel 2014). To subvert detection by extracellular immune receptors, many pathogens have evolved effector proteins that are secreted directly into the plant cell to either dampen plant immune responsiveness (Zhou et al. 2011) or produce metabolic environments favorable to pathogen growth (Chen 2014). In response to this effector secretion, plants have evolved a second, intracellular layer of pathogen recognition. This intracellular recognition is mediated by immune receptors called nucleotide-binding leucine-rich repeat proteins (NLR) (Flor 1971; Johal and Briggs 1992; Białas et al. 2021). Following pathogen perception, PRRs and NLRs mediate defense responses using partially overlapping mechanisms that result in the production of reactive oxygen species, regulation of phytohormones like salicylic and jasmonic acid, and increased expression of defense-related proteins, along with many other alterations (Klessig et al. 2000; Maximova et al. 2003; Kobayashi et al. 2012; Ngou et al. 2021).

NLR genes have received particular attention because of their ability to cause hypersensitive response, a type of qualitative resistance typified by programmed cell death and subsequent cessation of disease progression (Dangl and Jones 2001; Jones and Dangl 2006). NLR-mediated defense was originally discovered by Henry Harold Flor in the 1940s while breeding flax cultivars resistant to the rust pathogen *Melampsora lini* (Flor 1971). Flor originated the concept of gene-for-gene plant-pathogen interactions, wherein a specific NLR protein recognizes a specific pathogen effector, leading to resistance. Since then, much work has been done to characterize NLR-effector interactions. We now know of at least nine unique molecular mechanisms for NLR-mediated recognition of effector proteins (Kourelis and van der Hoorn 2018).

Most NLR genes contain just three domains. The center of the protein is composed of a nucleotide-binding (NB-ARC) domain that is homologous to human Apaf-1 and CED-4 NB-ARC domains (van der Biezen and Jones 1998). The N-terminal ends of NLRs are variable, containing either a Toll/interleukin-1 domain (TIR) or a coiled-coil motif (CC). And lastly, the C-terminal ends contain a set of leucine-rich repeats (LRR) that vary in length. Many NLRs maintain this canonical structure, but variation in gene architectures is widespread (Van de Weyer et al. 2019).

Due to their essential role in pathogen recognition and subsequent defense response, NLR genes and pathogen effectors are in a constant arms race. This tight co-evolutionary relationship has resulted in the expansion of NLR and effector repertoires (Meyers et al. 1998; Haas et al. 2009; Wang et al. 2021). For instance, it is not uncommon for NLRs to constitute 1-3% of a species’ total gene space (Zhang et al. 2016). At the same time, however, NLR copy number across plant genomes is also highly variable, from as few as 55 in watermelon (Lin et al. 2013) to as many as 2,151 in wheat (Andersen et al. 2020).

Likewise, limited evidence suggests NLR copy number can vary 1-2 fold within a species (Van de Weyer et al. 2019; Kim et al. 2021). Examining NLR complement across multiple individuals of the same species helps us understand how populations interact with their environments and provides insight into the ways NLR variation can be harnessed through breeding.

By virtue of being highly dynamic and repetitive, NLR genes are particularly difficult to assemble, annotate, and analyze. Genome resequencing efforts, moreover, are inherently reliant on reference genomes, limiting their utility for understanding natural variation in gene content, novel domain architectures, and genomic organization. Thus, genome-wide analysis of NLR variation across a species requires high quality *de novo* genome assemblies or the use of enrichment methods to perform high throughput sequencing of target genes (Jupe et al. 2013; Stam et al. 2016). Here, we explored the NLR content of 11 high quality genome assemblies from *Theobroma cacao,* the chocolate tree. We found NLR copy number was highly variable across genotypes, a phenomenon largely driven by tandem and segmental duplications. Our results provide additional insight into the evolution of NLRs within a species and suggests local duplications can drastically alter gene content over short evolutionary time scales.

## Materials and Methods

### Genome assembly and annotation

We analyzed 11 highly contiguous genomes, including both cacao reference genomes, Criollo B97-61/B2 v2.0 (GCA_000208745.2) and Matina 1-6 v2.1(GCA_000403535.1) (Motamayor et al. 2013; Argout et al. 2017), as well as nine other genomes that we sequenced, assembled, and annotated: CCN-51, GU-257E, ICS-1, NA-246, IMC-105, NA-807, Pound-7, SCA-6 (GCA_035896635.1), SPEC 54/1.

All assemblies were completed using Illumina 10X linked reads, collected and sequenced as previously described (Hämälä et al. 2021; Winters et al. 2024). Briefly, linked-reads were assembled using Supernova v2.1 (Weisenfeld et al. 2017) at five different raw read coverage depths: 56x, 62x, 68x, 75x, and 85x. The number of reads included in each coverage depth was determined according to estimated genome size. Each of the five coverage depths had two pseudohaplotype assemblies, one of which was chosen for post-processing. From the resulting five pseudohaplotype assemblies, a single representative was chosen as the meta-assembly backbone using a combination of metrics. These metrics included: completeness of benchmarking universal single copy orthologs (BUSCO) (Simão et al. 2015), contig and scaffold L50, and an assembly size that was consistent with the estimated haploid genome size. The remaining pseudohaplotype assemblies were then used to bridge gaps and join contigs, iteratively improving the meta-assembly backbone for each genotype. Assembly errors were corrected with TigMint (Jackman et al. 2018) before being re-scaffolded by ARCS (Yeo et al. 2018). Gaps were filled using GapFiller v1.10 (Boetzer and Pirovano 2012) and the resulting assembly for each genotype was called its meta-assembly. Chloroplast, mitochondria, and non-embryophyte contaminant sequences were removed using the BLAST-based procedure described previously (Winters et al. 2024). Finally, each meta-assembly was ordered and oriented onto pseudomolecules (chromosomes) using RaGOO (Alonge et al. 2019) and the *T. cacao* Matina 1-6 v1.1 genome assembly (Motamayor et al. 2013).

Before beginning annotation of the meta-assemblies, regions containing a high density of repeats or transposable elements were identified and masked using the MAKER-P repeat masking protocol (Campbell et al. 2014). A diverse set of tissues were sampled to generate transcripts that could be used as genome annotation evidence, and transcripts were created using the *de novo* assembly protocol described previously (Winters et al. 2024). Meta-assembly annotation took place in two steps. First, annotations from the Criollo B97-61/B2 v2.0 and Matina 1-6 v2.1 reference genomes were transferred to the meta-assemblies using the FLO pipeline (https://github.com/wurmlab/flo). Second, the assembled annotation evidence, as well as evidence from nine other species in Malvaceae, was used to create *de novo* annotations via the MAKER pipeline (Holt and Yandell 2011). These steps resulted in highly contiguous and complete genome assemblies that were suitable for analysis of complex, repeat rich genomic regions.

### Genotype phylogeny

We used benchmarking universal single-copy orthologs (BUSCO) to create a maximum likelihood phylogeny for the 11 genotypes analyzed in this study, as well as four non-cacao *Theobroma spp.* (Simão et al. 2015). First, we extracted complete BUSCOs from each genotype’s predicted proteome. Sequences for wild *Theobroma spp.* were extracted from the transcriptome assemblies described previously (Winters et al. 2024). Only complete BUSCOs present in 4 or more species were used for phylogenetic analysis. We then aligned each set of BUSCOs using MAFFT (L-INS-i) (Katoh et al. 2005) and constructed gene trees using FastTree v2.0 (Price et al. 2010). Lastly, we created a species tree from all 1,364 gene trees using the coalescent-based species tree estimation program ASTRAL (Mirarab et al. 2014).

We collected information on susceptibility phenotypes from the International Cacao Germplasm Database (http://www.icgd.rdg.ac.uk/) for three problematic cacao diseases: Ceratocystis wilt of cacao (CWC), frosty pod rot (FPR), and witches’ broom disease (WBD). These diseases were chosen because they had the largest collection of phenotype information in the database. Data on black pod rot phenotypes (*P. palmivora*) were taken from a previously published study (Fister et al. 2020). We filtered phenotype information using several criteria. First, we removed any information that was based on disease incidence in the field, since this is highly dependent on environmental conditions. Second, we removed sources of information that did not contain discernable phenotypes, e.g. “intermediate”. Lastly, since most clones have multiple phenotype estimates, we classified phenotype classes as numeric values and took the average, i.e. susceptible = 1, moderately susceptible = 2, moderately resistant = 3, and tolerant = resistant = 4. Phenotype averages that were ≤ 2 were considered susceptible and **≥** 3 were considered resistant.

### NLR classification and categorization

We identified NLR genes using a combination of custom, domain-specific hidden Markov models (HMM) and the NLR-identification tool NLR-parser (Steuernagel et al. 2015). We created a set of cacao-specific HMMs that were designed to detect homology with three canonical NLR domains: nucleotide-binding domain (NB-ARC), resistance to powdery mildew domain (RPW8), and the Toll/interleukin-1 receptor domain (TIR). In plants, each of these domains are diagnostic for the NLR gene family (Van de Weyer et al. 2019). HMMs were created by first identifying domains in each genome’s predicted proteome using Interproscan v5.32-71.0 (-appl Pfam) (Quevillon et al. 2005).

Proteins containing high confidence hits to NB-ARC (PF00931), TIR (PF01582 and PF13676), or RPW8 (PF05659) Pfam domains (e-values ≤ 1e-60 for NB-ARC or 1e-40 for TIR and RPW8) were used for further HMM construction (Mistry et al. 2021). This identified thousands of proteins containing NB-ARC and/or TIR domains. Domain sequences were then extracted from their respective proteins and aligned using MAFFT (--auto) (Katoh et al. 2005). After alignment, we constructed phylogenies from each domain alignment using RAxML (-m PROTCATJTT -p 1234). To limit the bias that highly similar, and therefore redundant, NB-ARC and TIR domain sequences could introduce during HMM classification, we used a clustering approach to identify unique sequences. This approach used USEARCH v11 (-cluster_tree -id 0.98) (Edgar 2010) to first cluster domain sequences that were ≥ 98% identical. A single representative was then selected from each cluster. This reduced the number of NB-ARC sequences by > 93% (152/2137) and the number of TIR sequences by nearly 50% (87/189). There were not enough RPW8 domains for redundancy to be an issue in HMM construction and classification, so no clustering and filtering was performed for this domain. Finally, representative NB-ARC, TIR, and RPW8 sequences were once again aligned using MAFFT (L-INS-i) and HMMs were built using HMMER v3.3 (hmmbuild) (Eddy 2011). This resulted in one HMM classifier for each of the three canonical domains.

The three HMM classifiers were then used to detect the presence of NB-ARC, TIR, or RPW8 domains from the predicted proteomes of all 11 genotypes. Proteins containing at least one high confidence (e-value ≤ 1e-4) hit were classified as NLRs and carried forward for further analysis. Because the HMM classifiers were constructed using domain sequences, they were not able to identify proteins containing coiled-coil (CC) motifs. To address this problem, we used the NLR identification tool NLR-parser. NLR-parser uses the MEME v4.9.1 suite (Bailey et al. 2009) to detect small stretches of sequence that are highly similar to a pre-defined set of NLR-specific motifs (Jupe et al. 2012), including CC motifs. Therefore, proteins identified as having CC motifs by NLR-parser were incorporated into our list of putative NLRs. Domain architectures for all putative NLRs were then assessed using Interproscan v5.32-71.0 (-appl Pfam, COILS) and used to categorize NLRs into TNL, RNL, CNL, or NL using previously outlined criteria (Van de Weyer et al. 2019). Proteins containing a TIR domain were categorized as TNL and those containing an RPW8 were categorized as RNL. Proteins containing an Rx N-terminal domain (PF18052), or an NLR-parser CC annotation *and* a CC annotation from COILS were categorized as CNL. And lastly, proteins containing an NB-ARC domain *and* a leucine-rich repeat (LRR) domain, but no other domains, were categorized as NL (Figure 1A).

**Figure 1:**
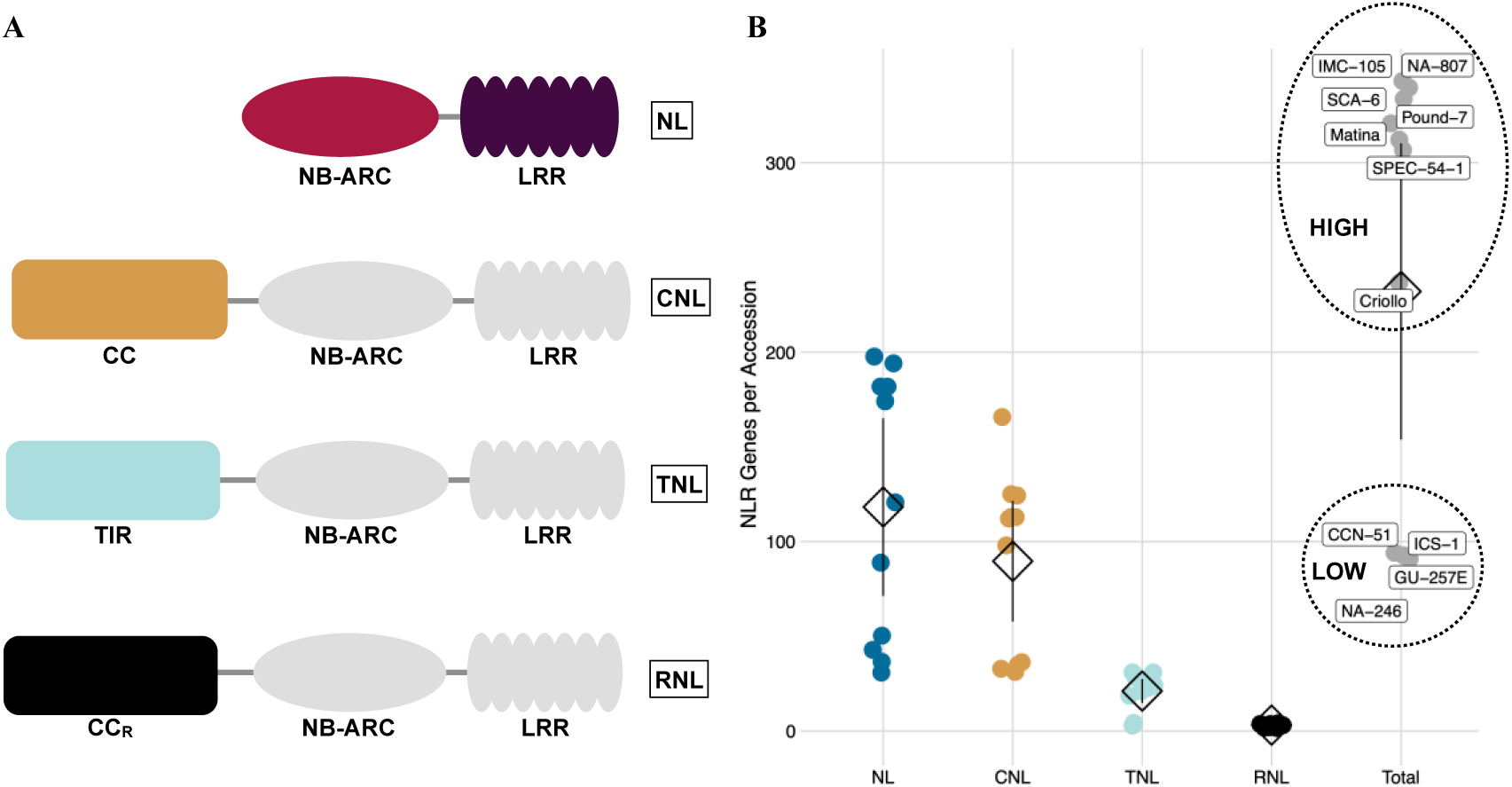
NLR architecture and copy number across cacao genomes. (A) The four canonical NLR architectures. NLR genes were classified as NL, CNL, or RNL according to their domain architecture. All NLRs contain an NB-ARC and LRR domain. This NB-ARC/LRR backbone either occurs in isolation (NL), or with one of three other domains, CC (CNL), TIR (TNL), or CC_R_ (RNL). (B) NLR copy number variation (CNV) across all classes and genotypes. NL, CNL, TNL, and RNLs are shown as blue, yellow, teal, and black, respectively. Each point represents the number of NLR copies for a particular genotype. Means are represented by diamonds. Lines represent 95% confidence intervals. High CNV genotypes had significantly more NLR genes than low CNV genotypes (mean difference = 225.11, Mann-Whitney test: p-value < 0.01). Other than the RNL class, differences in mean NLR number between Low CNV and High CNV genotypes were significant for all classes (negative binomial GLM: NLR # ∼ CNV Group + NLR Class + CNV Group * NLR Class, adjusted p-values < 0.01).

### Gene duplication analysis

Gene duplication histories were characterized using MCScanX’s duplicate gene classifier (Wang et al. 2012). First, we performed all-by-all BLASTp searches using each genotype’s predicted proteome with an e-value cutoff of 1e-10. These BLAST hits were then used as input to categorize duplication types. To do so, MCScanX first ordered genes according to their chromosomal location and categorized them as singletons. BLAST hits were then used to identify genes containing hits elsewhere in the genome. Any gene containing a hit elsewhere was called a dispersed duplicate. Dispersed duplicates were then further categorized as proximal duplicates if they were no more than 20 genes away from a BLAST hit, and tandem duplicates if they were one gene away from a BLAST hit. Lastly, genes classified as anchors by MCScsanX were categorized as segmental or whole genome duplicates (WGD).

NLR clustering was performed using a custom set of R and Bash scripts. First, we calculated the number of NLR genes in non-overlapping genomic windows of 1 Mbp. More than 50% of an NLR needed to overlap a window for it to be counted. Adjacent windows that both contained at least one NLR were then merged. This process was repeated until there were no remaining windows that could be merged. Sets of merged windows were considered NLR clusters if they contained ≥ 3 NLR genes.

### Genome synteny analysis

Pairwise comparisons of genome collinearity were performed using MCScanX (match_score = 50, gap_penalty = -1, match_size = 5, max_gaps = 20, repCut = 300, repDiv = 30). First, putative orthologs were identified with all-vs-all BLASTp (e-value 1e-10) and used to define collinear blocks according to the MCScanX algorithm. MCScanX accomplished this by first sorting BLAST hits according to their chromosomal positions. Long chains of collinear genes were then identified and collinear blocks longer than five genes were reported. Lastly, adjacent, collinear gene pairs were then used as anchors to align collinear blocks, identifying syntenic regions between genomes.

### Pseudogene identification

We identified pseudogenized NLR genes using the MAKER-P pseudogene identification pipeline (Zou et al. 2009; Campbell et al. 2014). First, we searched for genomic regions containing high sequence similarity to NLR genes using tBLASTn (e-value ≤ 1e-20). These regions of sequence similarity were then used as input into the pseudogene pipeline. Non-genic regions were sorted to remove hits ≤ 30 amino acids in length and ≤ 40% identity. Regions passing these filters were considered putative pseudoexons.

Putative pseudoexons that significantly matched (e-value < 1e-5) repetitive sequences, as defined in RepBase v.12 (Jurka et al. 2005), were removed. We linked the remaining pseudoexons together to form contigs based on two criteria: (1) the best BLAST hit for both pseudoexons was the same parent NLR, and (2) the sequence space between the matching pseudoexons was inside the 99^th^ percentile of the intron length distribution. These contigs represented putative pseudogenes. NLR integrated domains, i.e. non-canonical NLR domains that are fused to NLR genes, presented a challenge because they would result in the identification of non-NLR pseudoexons and subsequent pseudogenes. Therefore, putative pseudogenes were translated in all six frames and their domains were identified using Interproscan v5.32-71.0 (-appl Pfam). Pseudogenes that did not contain any common NLR domains were removed. We considered the following NLR domains as common: LRR (PF00560, PF07725, PF12799, PF13855), NB-ARC (PF00931), TIR (PF01582, PF13676), RPW8 (PF05659), and CC (PF18052). This filtered set of pseudogenes represented putatively non-functional NLR genes.

### Transposable element analysis

We annotated transposable elements (TE) from each genome according to the MAKER-P repeat masking protocol, as previously described (Winters et al. 2024). First, miniature inverted-repeat transposable elements (MITE) were identified using MITE-Hunter (Han and Wessler 2010). Likewise, long terminal repeat retrotransposons (LTR) were identified using LTRharvest/LTRdigest (Ellinghaus et al. 2008; Steinbiss et al. 2009). *De novo* repetitive sequences were predicted using RepeatModeler1 (http://www.repeatmasker.org/RepeatModeler). Predicted TEs were then searched against a SwissProt and RefSeq protein database. Any TEs containing significant hits to the database were excluded from further analysis. We chose to analyze five TE classes: DNA transposons, LINE, SINE, and LTR retrotransposons, and rolling circle Helitrons.

### Statistical analyses

We performed all statistical analyses using R v3.6 (Team R.C. 2013). Negative binomial regressions were performed using the MASS v7.3-53.1 package (Venables and Ripley 2002) and pairwise significance was calculated using the emmeans v1.5.4 package (Lenth et al. 2018). We checked all model assumptions using performance v0.8.0 (Lüdecke et al. 2021). All confidence intervals were calculated using Hmisc v4.4-2 (Harrell and Dupont 2006). Plots were created using ggplot2 v3.3.5 (Wickham 2009) and genoPlotR v0.8.11 (Guy et al. 2010).

## Results

### Cacao genotypes displayed extensive copy number variation in NLR genes

We used 11 genotypes for this study, each belonging to one of seven previously described (Motamayor et al. 2008) genetic groups displaying divergence by distance: Criollo B97-61/B2 (Criollo), Matina 1-6 v2.1 (Amelonado), CCN-51 (Hybrid), GU-257E (Guiana), ICS-1 (Hybrid), NA-246 (Marañón), IMC-105 (Iquitos), NA-807 (Nanay), Pound-7 (Nanay), SCA-6 (Contamana), and SPEC 54/1 (Iquitos). The exceptions to this were CCN-51 (Iquitos x Amelonado x Criollo) and ICS-1 (Amelonado x Criollo), both of which are hybrids. In total, we identified 2,563 NLR genes across 11 genomes (Table S1). We further categorized these NLRs into groups based on their domain architecture (Figure 1A).

Typical NLR genes are tripartite, possessing a variable N-terminal domain, a conserved NB-ARC domain, and a C-terminal end containing a set of leucine-rich repeats of varying length. Based on the presence and/or absence of these domains and their organization, we categorized NLR genes into four classes: NL, CNL, TNL, and RNL (Figure 1A). RNL copy number (Mean_RNL_ = 3.09, SEM_RNL_ = 0.25) appeared to be low and conserved across genotypes, consistent with their role as an ancient clade of helper NLRs (Figure 1B) (Jubic et al. 2019; Lapin et al. 2019). Copy number in NL (Mean_NL_ = 118.27, SEM_NL_ = 21.05), CNL (Mean_CNL_ = 89.64, SEM_CNL_ = 14.28), and TNL classes (Mean_TNL_ = 21.09, SEM_TNL_ = 2.86), however, was highly variable, with particular divergence seen in both NL and CNL. Total NLR number varied across genotypes, hereafter referred to as copy number variation (CNV), and fell into two discrete classes: genotypes with high NLR copy number (Mean_HighCNV_ = 314.86, SEM_HighCNV_ = 13.72) and genotypes with low copy number (Mean_LowCNV_ = 89.75, SEM_LowCNV_ = 2.98), hereafter referred to as High CNV and Low CNV, respectively (Figure 1B). There was a 3-fold difference in NLR copy number between these two groups (mean difference = 225.11, Mann-Whitney test: p-value < 0.01), most of which was driven by expansion and/or contraction of the NL and CNL classes (negative binomial GLM: NLR # ∼ CNV Group + NLR Class + CNV Group * NLR Class, adjusted p-values < 0.01).

### NLR copy number variation was not distributed evenly throughout the genome

NLR genes were distributed across all 10 cacao chromosomes, but most (45-84%) were localized to just four: Chr5, Chr6, Chr7, and Chr10 (Figure 2A). This was true for both Low CNV and High CNV genotypes. Consistent with a birth-and-death model, these four chromosomes also contained the greatest concentration of pseudogenes in the High CNV genotypes (Figure 2B). Low CNV genotypes, however, were more variable. Three of the four Low CNV genotypes, CCN-51, GU-257E, and NA-246, had zero NLR pseudogenes, while ICS-1 had 2x the number of NLR pseudogenes as it did NLRs (Figure 2B). ICS-1 pseudogenes were distributed across chromosomes 5, 6, 7, and 10, similar to the High CNV genotypes, but patterns of gene duplication varied. While the high CNV genotypes had approximately 1.6 pseudogenes for every 1 parent NLR (Mean = 1.63, SEM = 0.09), ICS-1 had 4.6. That is, ICS-1 had 198 pseudogenes derived from just 43 parent NLRs. These results indicate NLR genes, at least in the High CNV genotypes, expanded on a small number of chromosomes, helping drive the observed differences in copy number across genotypes. This NLR expansion was then followed by pseudogenization according to a birth-and-death model. This process, however, likely occurred differently in ICS-1, for reasons we elaborate in the discussion.

**Figure 2:**
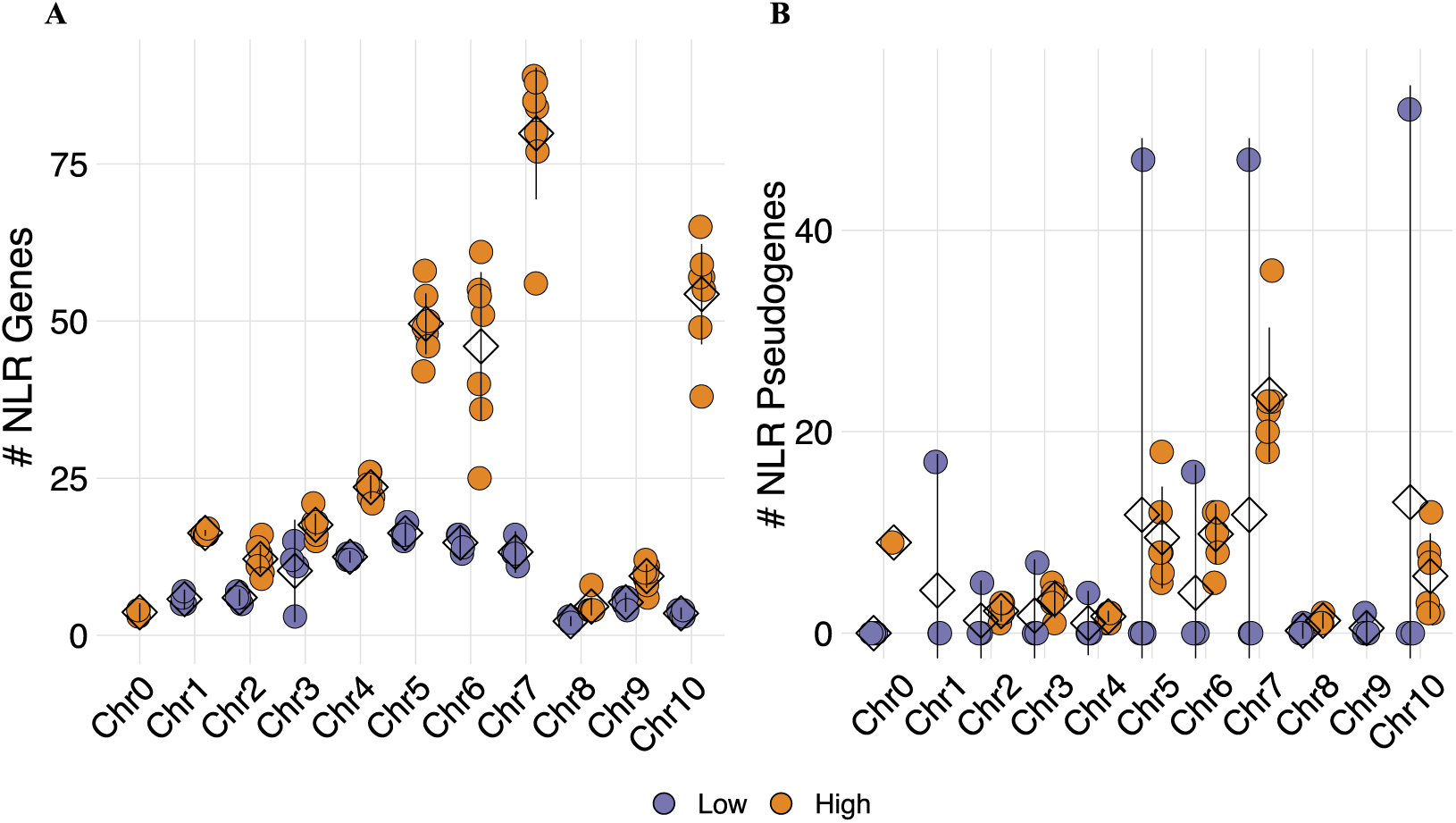
Distribution of NLR genes across each genome. (A-B) Number of NLR genes or NLR pseudogenes on each chromosome. Orange depicts High CNV genotypes and purple depicts Low CNV genotypes. Each point represents the number of NLR genes or NLR pseudogenes for a particular genotype. NLR genes or NLR pseudogenes on Chr0 do not belong to one of the 10 chromosome-oriented scaffolds. Means are represented by diamonds. Lines represent 95% confidence intervals. Where the lower tail of the confidence interval is negative, the line is truncated at zero. If there is no variance for an observation, no confidence interval is shown.

### High and low copy number genotypes evolved independently multiple times

To test whether low versus high copy number was segregating across the cacao phylogeny, we created a species tree from 1,364 single copy ortholog trees (Figure 3). Population-level relationships were recovered for Nanay but not for Iquitos. This is consistent with other phylogenetic trees (Winters et al. 2024), and likely occurred because SPEC 54/1 is highly differentiated from other Iquitos genotypes. While low versus high copy number was consistent within each clade, e.g. both NA-807 and Pound-7 belong to High CNV, there was no consistency within populations, e.g. SPEC 54/1 versus IMC-105.

**Figure 3:**
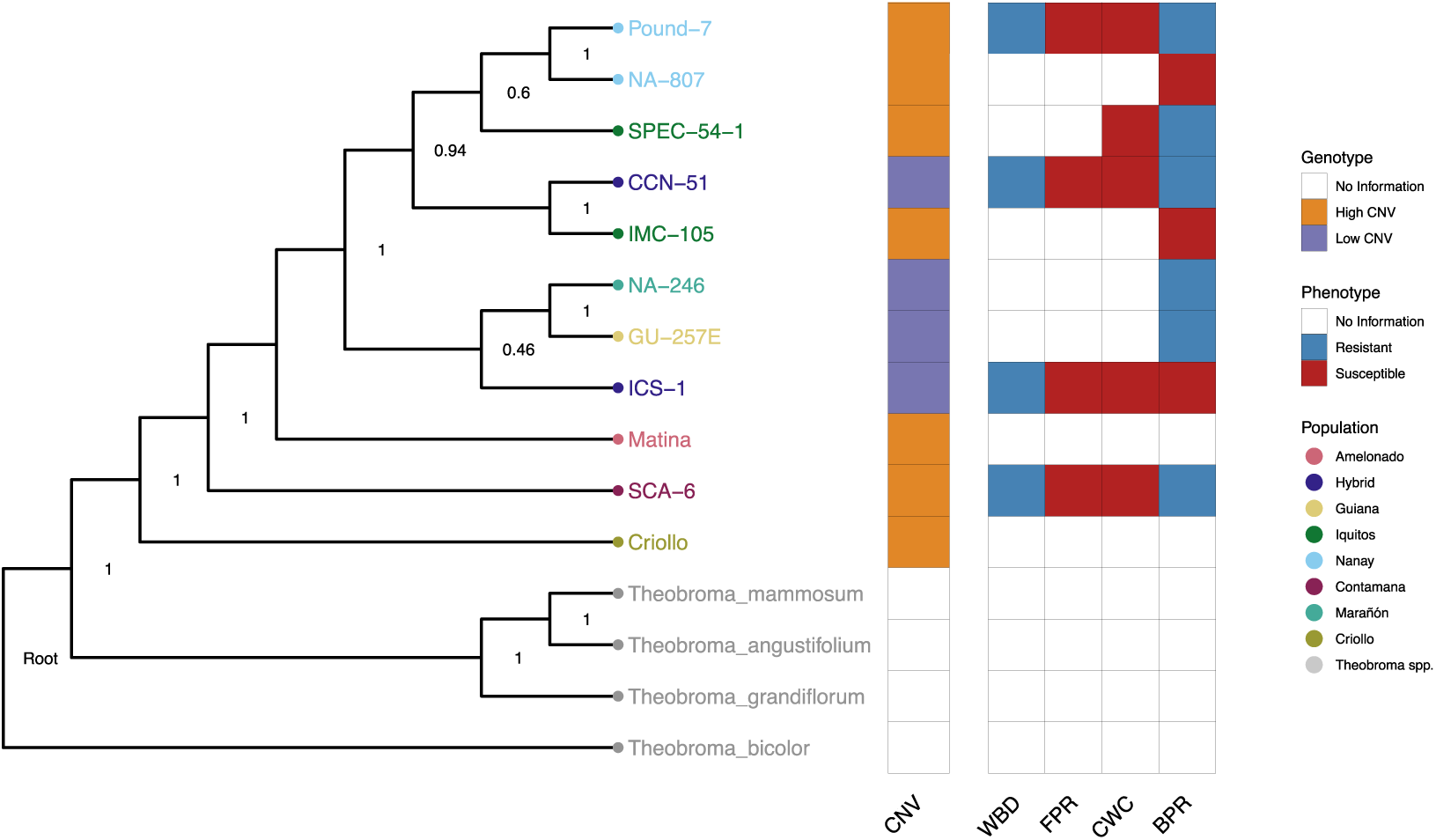
Phylogeny of cacao genotypes sampled for this study. Phylogenetic tree of the 11 cacao genotypes used in this study, constructed using 1,364 single copy genes. Four non-cacao species of *Theobroma* were additionally used as outgroups. Numbers on each node represent posterior probability support values calculated by ASTRAL. With the exception of CCN-51 and ICS-1, both of which are hybrids, tip colors indicate population membership. Non-cacao *Theobroma spp.* are shown in grey. CNV class (High, Low, or No Information) of each genotype is shown in orange, purple, or white, respectively. Disease phenotypes are shown for witches’ broom disease (WBD), frosty pod rot (FPR), *Ceratocystis* wilt of cacao (CWC) and black pod rot (BPR). Blue indicates resistant, red indicates susceptible, and white indicates no information was available.<colcnt=1>

Likewise, more broadly defined clades are also inconsistent with respect to their CNV group, e.g. GU-257E, NA-246, and SCA-6. These results indicate High CNV, as a trait, evolved independently multiple times. It is likely, however, that, given a larger sample size, copy number variation among genotypes would become less discrete, forming a more continuous distribution that fills in the gap between High and Low CNV genotypes. Thus, it may be necessary to assess NLR copy number variation across the phylogeny as quantitative rather than qualitative. Lastly, CNV group was not associated with resistance to Ceratocystis, black pod rot, frosty pod rot, or witch’s broom disease (Table S2, Figure 3).

### Annotation quality varied across genotypes but did not explain NLR copy number variation

To determine whether differences in total annotated gene number across genomes drove variation in NLR copy number, we analyzed three separate metrics of annotation quality for each genome (Figure 4). We began by investigating differences in total annotated gene space among Low CNV and High CNV genotypes (Table S4, Figure 4A). While the Low CNV genotypes possessed fewer annotated genes (Mean_Gene #_ = 25,064.25 genes) than High CNV genotypes (Mean_Gene #_ = 26,145.14 genes), the difference was not significant (mean difference = 1080.89 genes, Mann-Whitney test: p-value > 0.05). Moreover, this approximately 4% variation in total gene content between High and Low CNV genotypes is much lower than the approximately 300% variation in NLR copy number. Thus, the only way differences in annotated gene content could have driven the observed patterns of NLR copy number variation is if annotation was systematically biased against NLRs. Next, we used MAKER’s annotation edit distance (AED) to assess the quality of annotated NLR genes. AED is a measure of how well annotations match aligned transcripts and protein data and has scores ranging from 0 to 1. An AED of 0 indicates a perfect match between an annotation and its evidence, while an AED of 1 indicates complete discordance.

**Figure 4:**
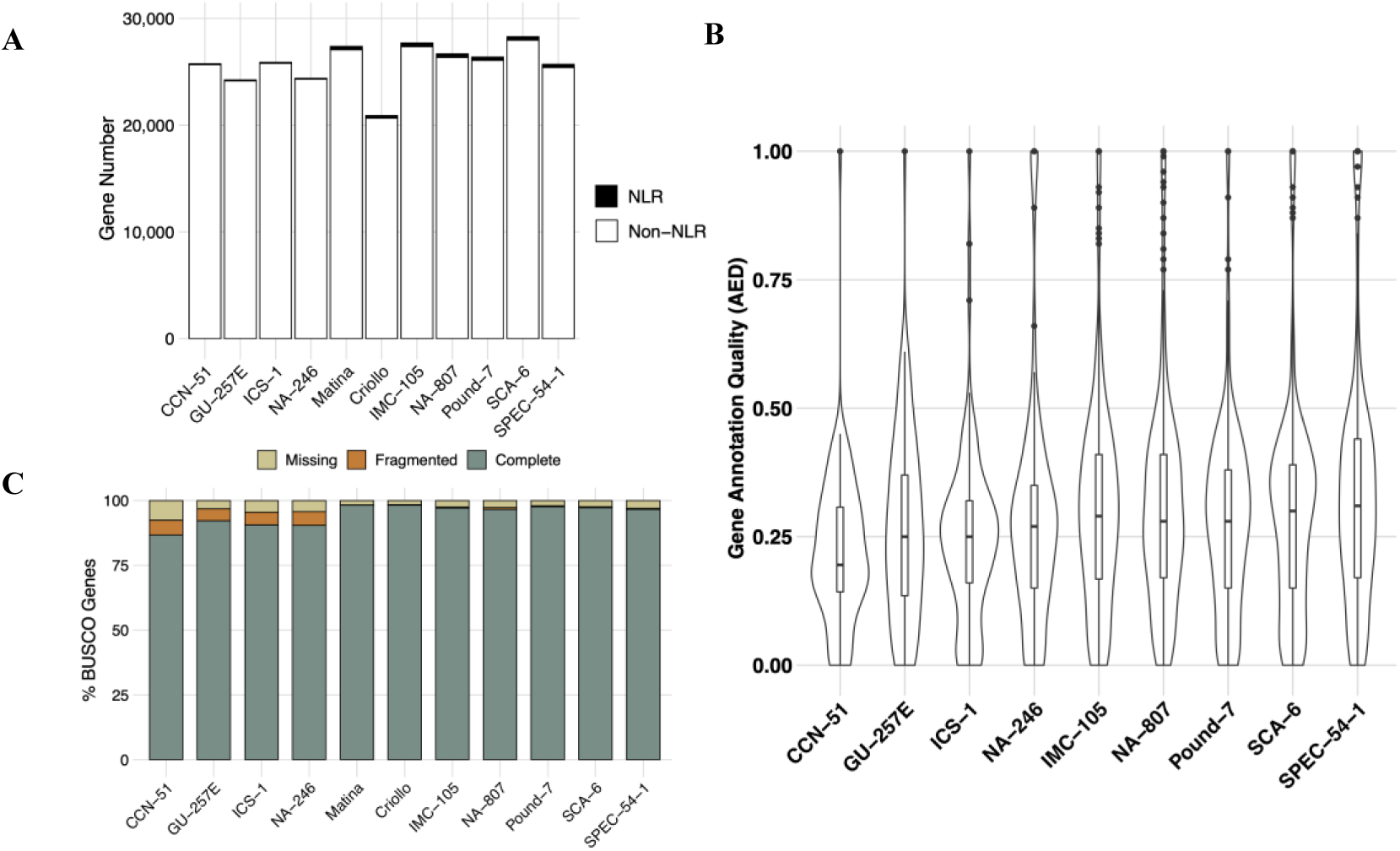
Genome annotation quality metrics. (A) The total number of genes annotated in each of the 11 genomes used in this study, separated by Low CNV (left) and High CNV (right). NLR genes are shown in black and non-NLR genes are shown in white. There was no significant difference in gene number between Low CNV and High CNV genotypes (mean difference = 1080.89 genes, Mann-Whitney test: p-value > 0.05). (B) Distribution of AED scores for each genotype’s classified NLR genes. Mean AED score was not significantly different between Low CNV and High CNV groups (mean difference = 0.018, t-test: p-value < 0.001). (C) BUSCO completeness for each genome used in this study, separated by Low CNV (left) and High CNV (right). The proportion of complete, fragmented, and missing BUSCOs are shown in green, orange, and beige, respectively. Differences in the mean proportion of complete, fragmented, and missing genes between Low CNV and High CNV genotypes were significant (one-way ANOVA: Proportion ∼ CNV Group + BUSCO Class + CNV Group * BUSCO Class, p-value < 0.001; Tukey’s HSD, adjusted p-value < 0.01).

Annotations with AED scores < 0.2 are considered extremely high quality (Holt and Yandell 2011). Mean AED was significantly different between Low CNV genotypes and High CNV genotypes (t-test: p-value < 0.001; Figure 4B). However, both CNV groups had AED distributions with means centered near 0.2 (0.19 for Low CNV and 0.20 for High CNV, mean difference = 0.018). This indicates NLR annotations for both Low CNV and High CNV genotypes are not likely to be spurious. Lastly, we assessed BUSCO completeness across all 11 genotypes (Table S3). The Low CNV group had a significantly lower proportion of both complete and fragmented BUSCOs relative to the High CNV group (one-way ANOVA: Proportion ∼ CNV Group + BUSCO Class + CNV Group * BUSCO Class, p-value < 0.001), indicative of less complete genome annotations (Figure 4C). The mean difference for complete and fragmented BUSCOs was 7.38% and 4.75%, respectively. The average proportion of complete BUSCOs for both CNV groups, however, was **≥** 90%, which is generally considered highly complete (Simão et al. 2015). Together, these results suggest differences in NLR copy number are not the result of technical differences in annotation quality but are due to authentic differences in duplication histories across genotypes.

### Variation in transposable element content did not explain NLR copy number variation

Gene duplication is a common feature of plant genomes. Duplications can take place through many mechanisms, such as whole genome duplication (Jiao et al. 2011; One Thousand Plant Transcriptomes Initiative 2019), unequal crossing over during meiosis (Jelesko et al. 1999), non-homologous end joining during DNA damage repair (Vaughn and Bennetzen 2014), slipped-strand mispairing (Xu et al. 2021), and transposable element-mediated operations (Kim et al. 2017). The last of these possibilities was recently shown to play a role in NLR duplication and diversification in two species in the genus *Capsicum*: *C. baccatum* and *C. chinense* (Kim et al. 2017). To investigate whether TEs were responsible for the patterns of NLR copy number variation across cacao genotypes, we annotated both well-known and uncharacterized TEs from 10 of the 11 genotypes in this study (Supplemental Table S5). We examined the abundance of five known TE classes: DNA transposons (Kapitonov and Jurka 2006), SINE, LINE, and LTR retrotransposons (Xiao et al. 2008; Kramerov and Vassetzky 2011), and rolling circle Helitrons (Kapitonov and Jurka 2001). LTR elements were by far the most abundant, followed by DNA, LINE, SINE, and Helitron elements, respectively (Figure 5). There were no significant differences in TE abundance between High CNV and Low CNV genotypes (negative binomial GLM: # TE ∼ CNV + TE Class + CNV*TE Class, adjusted p-values > 0.05; Figure 5).

**Figure 5:**
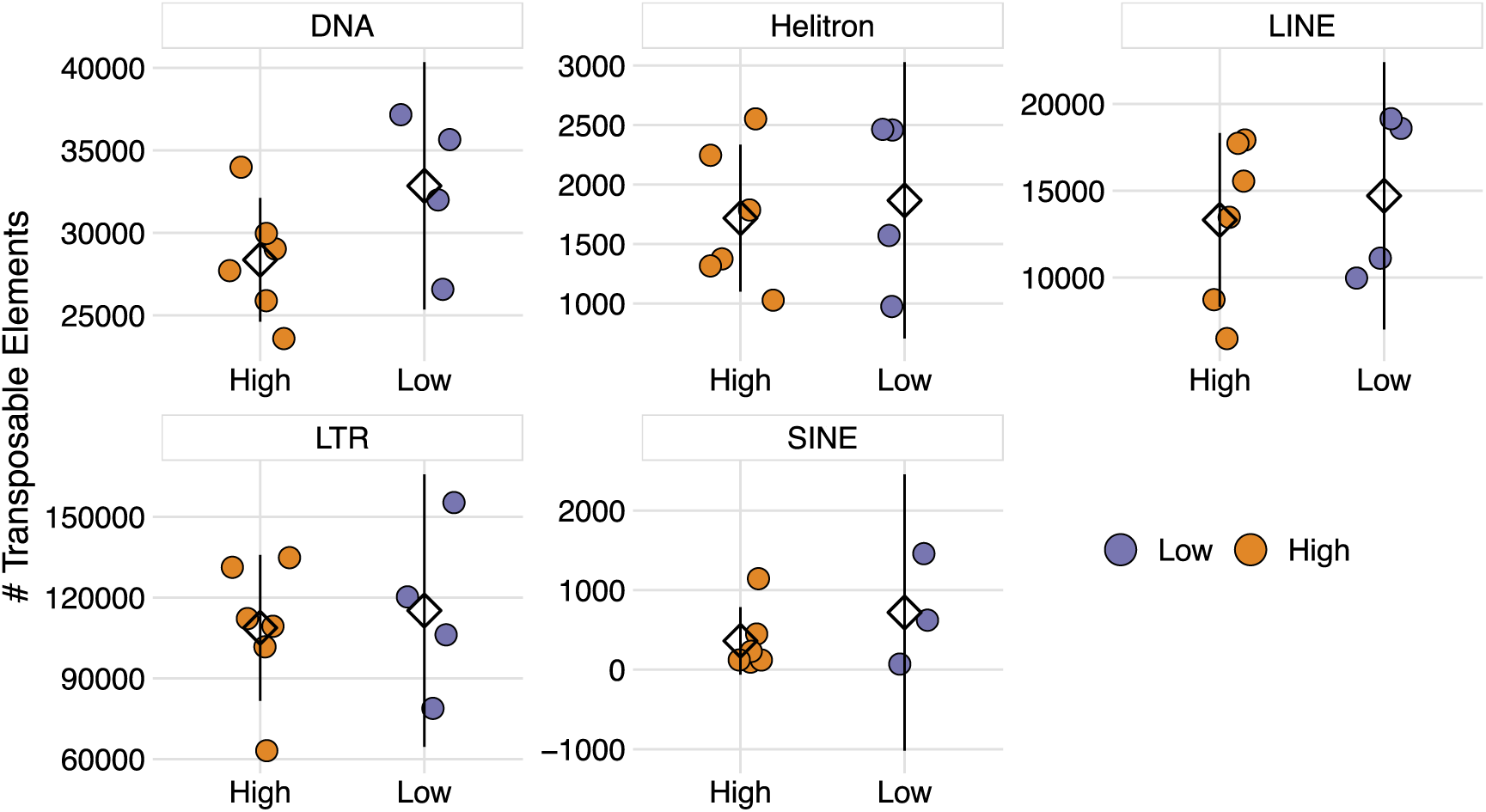
Transposable element abundance for High and Low CNV genotypes. Abundance of the five most common transposable elements in cacao genomes. Orange depicts High CNV genotypes and purple depicts Low CNV genotypes. Each point represents the number of transposable elements for a particular genotype. Means are represented by diamonds. Lines represent 95% confidence intervals. Differences in mean TE abundance between Low and High CNV genotypes were not significant (negative binomial GLM: # TE ∼ CNV + TE Class + CNV*TE Class, adjusted p-values > 0.05).

Aggregate patterns of TE abundance across genomes, however, may mask differences in local TE density that drive gene duplications. Therefore, we also investigated the abundance of TE classes on each chromosome for both Low and High CNV genotypes. Patterns of TE density across chromosomes did not match those seen for NLRs (Figure 6). That is, very few TE classes had significantly higher abundance among NLR-dense chromosomes, i.e. Chr5, Chr6, Chr7, and Chr10 (negative binomial GLM: # TE ∼ Chrom + TE Class + Chrom*TE Class). There were significantly more Helitron TEs on Chr5, Chr6, and Chr10 relative to Chr8 (adjusted p-values < 0.05), and significantly more LTR TEs on Chr5 relative to Chr8 (adjusted p-value < 0.01). This, however, is primarily because Chr8 had the lowest TE abundance across all classes, and likely has no influence on NLR accumulation. Moreover, none of the chromosomes displayed significant differences in TE content between Low and High CNV genotypes (negative binomial GLM: # TE ∼ CNV + Chrom + TE Class + Chrom*TE Class*CNV, adjusted p-values > 0.05). These results suggest variation in NLR content, both within and between genomes, was not explained by variation in TE content and potential TE-mediated duplications.

**Figure 6:**
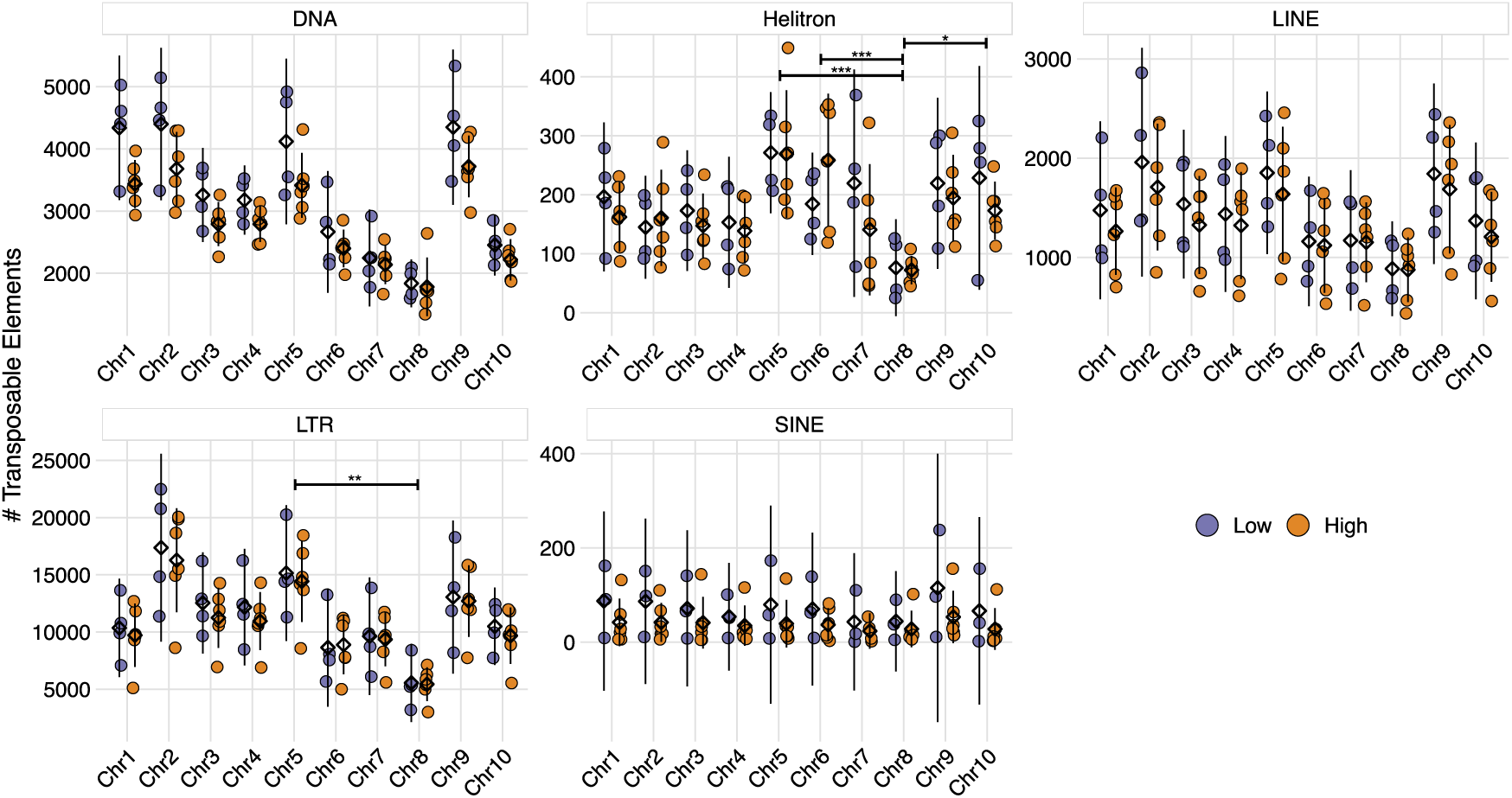
Density of the five most common transposable elements across each cacao chromosome. Orange depicts High CNV genotypes and purple depicts Low CNV genotypes. Each point represents the number TEs for a particular genotype. Means are represented by diamonds. Lines represent 95% confidence intervals. Stars indicate significant differences in mean TE abundance between chromosomes (negative binomial GLM: # TE ∼ Chrom + TE Class + Chrom*TE Class, adjusted p-values < 0.05). Differences in mean TE abundance between Low and High CNV genotypes on each chromosome were not significant (negative binomial GLM: # TE ∼ CNV + Chrom + TE Class + Chrom*TE Class*CNV, adjusted p-values > 0.05).

### Tandem and proximal duplications were primarily responsible for NLR copy number variation

Confident that differences in NLR copy number across genotypes were not due to technical discrepancies in annotation quality or mediated by transposable elements, we began investigating gene duplication histories across all 11 genotypes (Table S5). To do this, we used MCScanX’s duplicate gene classifier to categorize NLR genes as singletons, dispersed, proximal, tandem, or WGD/segmental duplicates (Figure S1). All NLRs were first classified as singletons, i.e. genes with no history of recent duplication (Figure S1A). If NLR genes contained significant BLAST hits elsewhere in the genome, they were reclassified as dispersed duplicates (Figure S1B). Dispersed duplicates were then further categorized as proximal or tandem based on distance between hits. If < 20 genes separated the NLR duplicates, they were considered proximal (Figure S1C). If NLR duplicates were immediately adjacent to one another, they were considered tandem (Figure S1D). Lastly, NLR duplicates that were anchors of collinear blocks, as defined by MCScanX’s algorithm (Wang et al. 2012), were classified as WGD/segmental duplicates (Figure S1E).

NLR genes had disproportionately higher tandem (mean difference = 23.9%) and proximal (mean difference = 24.3%) duplication rates relative to non-NLR genes (one-way ANOVA: Proportion ∼ Gene Type + Duplicate Type + Gene Type * Duplicate Type, p-value < 0.001; Tukey’s HSD, adjusted p-values < 0.001; Figure 7A). Likewise, NLR genes had a significantly lower proportion of both dispersed (mean difference = 7.1%) and segmental (mean difference = 17.5%) duplications relative to non-NLR genes (one-way ANOVA: Proportion ∼ Gene Type + Duplicate Type + Gene Type * Duplicate Type, p-value < 0.001; Tukey’s HSD, adjusted p-values < 0.01; Figure 7A). These results are consistent with previous findings that NLR evolution is primarily driven by local duplication events (Meyers et al. 1998; Meyers et al. 2003). There were very few singleton NLRs relative to non-NLR genes. Therefore, we focused the remaining analyses on tandem, proximal, dispersed, and segmental duplicates.

**Figure 7:**
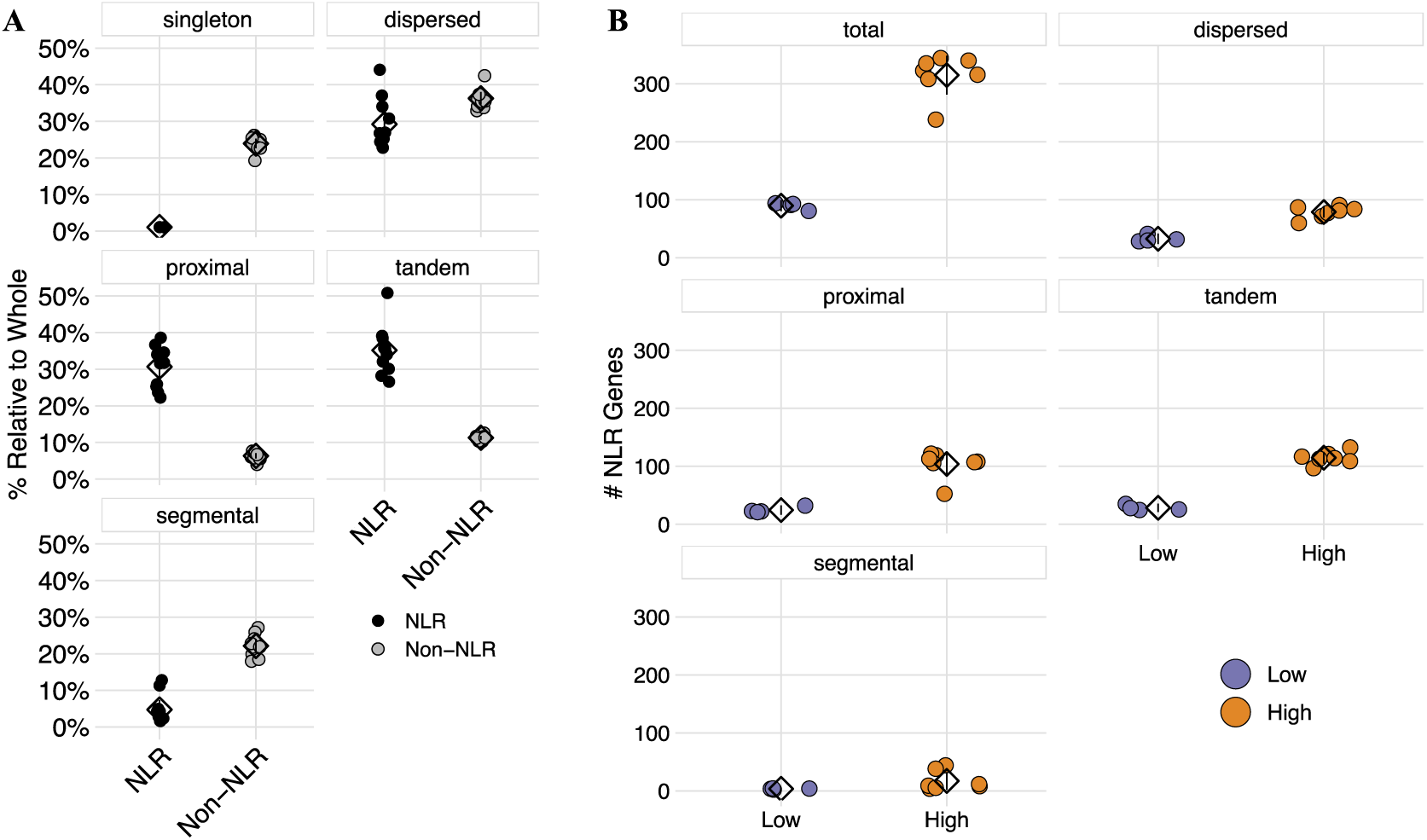
Patterns of NLR duplication across each genome. (A) The proportion of NLR (black) and non-NLR (grey) genes in each duplication class. Each point represents the proportion of NLR or non-NLR genes for a particular genotype. Means are represented by diamonds. Lines represent 95% confidence intervals. All differences in mean proportion between NLR and non-NLR genes were significant (one-way ANOVA: Proportion ∼ Gene Type + Duplicate Type + Gene Type * Duplicate Type, p-value < 0.001; Tukey’s HSD, adjusted p-values < 0.001). (B) The number of NLR genes belonging to each duplication class, for both Low CNV (purple) and High CNV (orange) genotypes. Points represent the number of NLR genes for a particular genotype. Means are represented by diamonds. Lines represent 95% confidence intervals. All differences in mean NLR number between Low CNV and High CNV groups were significant (negative binomial GLM: # NLR Duplicates ∼ Duplicate Type + CNV Group + Duplicate Type * CNV Group, adjusted p-values < 0.001).

While tandem and proximal duplications drove NLR evolution broadly, we next sought to determine how differences in NLR copy number arose across genotypes. To test this, we examined the NLR duplication history of both Low CNV and High CNV groups (Figure 7B). High CNV genotypes had significantly more NLR duplicates across all types (negative binomial GLM: # NLR Duplicates ∼ Duplicate Type + CNV Group + Duplicate Type * CNV Group, adjusted p-values < 0.001). Again, however, the largest difference was seen for tandem and proximal duplicates, which together accounted for 73.7% of the total difference in NLR number between High CNV and Low CNV genotypes (165.9/225.1). Thus, it appears that tandem and proximal duplication events not only drive differences between NLR and non-NLR genes, but also drive variation in NLR content across genotypes.

To determine whether duplication type was biased towards specific gene architectures, we calculated the number of NL, CNL, TNL, and RNL genes belonging to each duplication type. Across all four types of duplication, the proportion of NLRs in each architecture class was constant (Figure 8). There was, however, a large difference in NLR copy number within both NL and CNL. Thus, duplications, as a whole, were biased towards NL and CNL architectures, and this bias was consistent across duplication types.

**Figure 8:**
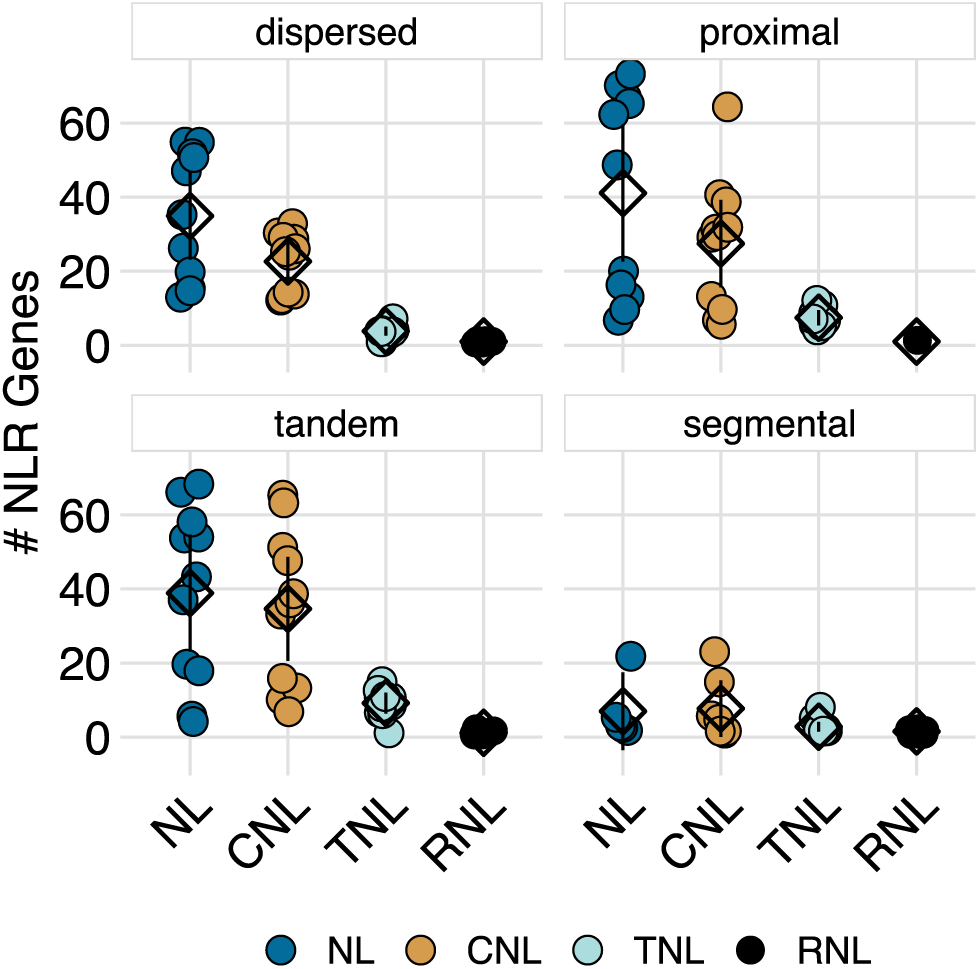
NLR duplications across domain architectures. NL, CNL, TNL, and RNLs are shown as blue, yellow, teal, and black, respectively. Each point represents the number of NLR copies for a particular genotype. Means are represented by diamonds. Lines represent 95% confidence intervals.

Most NLR copy number variation occurred on just four chromosomes: Chr5, Chr6, Chr7, and Chr10. To test whether tandem and proximal duplications were responsible for the formation of these NLR hotspots, we investigated the distribution of NLRs across both chromosomes and duplication types (Figure 9). While Chr7 harbors a higher number of dispersed duplicates than other chromosomes, NLR density on Chr5, Chr6, Chr7, and Chr10 was primarily driven by tandem and proximal duplications (Figure 9A). Likewise, differences in copy number between Low CNV and High CNV genotypes was associated with tandem and proximal duplications on these chromosomes. Consistent with these results, there was a significant difference in both the number and size of NLR clusters between Low CNV and High CNV genotypes (Figure 9B), a difference predominantly caused by proximal and tandem duplications (cluster number: mean difference = 6.32, Mann-Whitney test, p-value < 0.01; cluster size: mean difference = 11.48, Mann-Whitney test, p-value < 0.05). Thus, it appears that most NLR copy number variation is occurring on just four chromosomes, and these differences are principally due to proximal and tandem duplications driving expansion of NLR clusters. The expansion of one such NLR cluster can be seen in Figure S2. While the Low CNV genotypes have NLRs in this 2 Mbp region, cluster expansion via tandem and proximal duplications in High CNV genotypes is striking.

**Figure 9:**
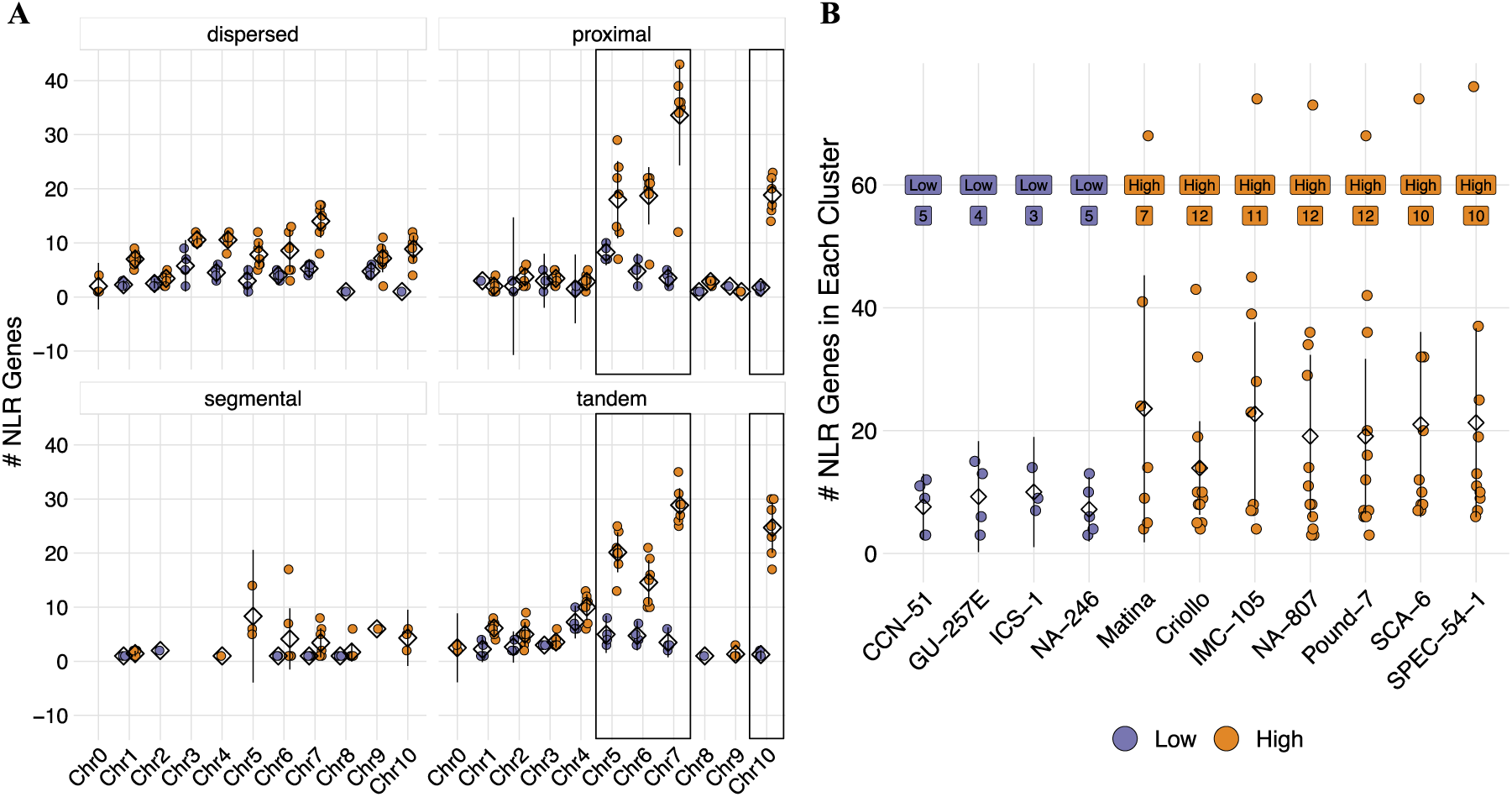
Number, size, and location of NLR clusters. (A) The genomic distribution of NLRs in each duplicate type, for Low CNV (purple) and High CNV (orange) genotypes. Each point represents the number of NLRs for a particular genotype. Boxes outline the four chromosomes with the highest NLR density. Means are represented by diamonds. Lines represent 95% confidence intervals. (B) Number and size of NLR clusters for each genotype, for Low CNV (purple) and High CNV (orange) genotypes. Each point represents a single NLR cluster. Mean cluster size for each genotype is represented by a diamond. Lines represent 95% confidence intervals. Boxed values indicate the number of NLR clusters for each genotype. Differences in both mean cluster number and mean cluster size between Low CNV and High CNV genotypes were significant (cluster number: mean difference = 6.32, Mann-Whitney test, p-value < 0.01; cluster size: mean difference = 11.48, Mann-Whitney test, p-value < 0.05).

## Discussion

Pathogen recognition is the first step in plant defense response and subsequent resistance or susceptibility. This recognition takes place either extracellularly, by PRRs, or intracellularly, by NLRs (Johal and Briggs 1992; Gómez-Gómez and Boller 2000). Co-evolution between pathogen effectors and plant NLR genes has resulted in an arms race typified by large and diverse repertoires of both effector and NLR gene families (Meyers et al. 1998; Haas et al. 2009; Wang et al. 2021). The large number of high-quality genomes sequenced over the last two decades have revealed huge variation in NLR copy number between even closely related species (Zhang et al. 2016). However, due to the high cost of sequencing, assembling, and annotating high-quality genomes, we still have a very limited understanding of genome-wide NLR diversity within a single species. In this study, we investigated the evolution of NLR content and location across 11 high-quality genome assemblies of the chocolate tree, *Theobroma cacao,* with the goal of gaining further insight into both NLR evolution and cacao’s interactions with its microbial environment.

NLR copy number was divided into two discrete groups, between which there was a 3-fold difference in gene content (Figure 1B). It is hard to know how this compares to other species, but the limited evidence we have suggests this is a high degree of variation. For instance, across 64 genotypes of *A. thaliana* Van de Weyer et al. found an approximately 1.5-fold variation in NLR copy number (Van de Weyer et al. 2019). Likewise, NLR copy number varied < 1-fold across five genotypes of *C. annuum* (Kim et al. 2017). Variation in NLR copy number across cacao genotypes may have many explanations, among them ancestral polyploidy, gene flow among populations, and divergence time. Polyploid and whole genome duplication events have played significant role in the speciation and diversification of land plants (Mandáková and Lysak 2018). Polyploid events are often followed by rapid diploidization, wherein duplicated genes gain new functions or are lost through pseudogenization (Scannell et al. 2006; Scannell et al. 2007), a process that can contribute to variation in gene content across a lineage (Mandáková and Lysak 2018). Cacao does not have an evolutionary history of polyploidy since the gamma triplication shared by all core eudicots (Argout et al. 2011; Jiao et al. 2011), thus whole genome duplication followed by differential loss of duplicated genes is unlikely to result in the observed pattern of NLR copy number variation. Limited gene flow between populations of cacao means variation is not homogenized through mating, allowing for greater diversification (Slatkin 1993). Long divergence times, i.e. time since speciation, means there is greater opportunity for variants to arise and fix in populations.

Cacao diverged from its most recent common ancestor 9.9 million years ago (Richardson et al. 2015). *A. thaliana,* in contrast, diverged from other *Arabidopsis* species approximately 6 million years ago (Novikova et al. 2016), and *C. annuum* diverged from other *Capsicum* species sometime in the last 3.4 million years (Särkinen et al. 2013; Carrizo García et al. 2016). While speciation time is certainly not the only factor controlling genetic variability, it is still important for population-level differentiation (Haag et al. 2005). Together, long divergence times and stratified populations may have helped facilitate large differences in NLR gene content across cacao genotypes. Selective forces driving the maintenance of high NLR copy number, however, are far less clear. NLR genes are energetically costly to produce in the absence of their cognate pathogen (Tian et al. 2003), and their mis-regulation can lead to autoimmunity (Lolle et al. 2017).Conversely, within-species NLR diversity is important for the colonization of new habitats in wild tomato populations (Stam, Silva-Arias, et al. 2019), and decreased NLR polymorphism is associated with greater conspecific negative density dependence (Marden et al. 2017). Whether variation in NLR copy number is beneficial or detrimental to plant fitness is therefore context dependent.

Differences in NLR copy number were not associated with resistance or susceptibility to a range of cacao diseases (Figure 3). This is largely unsurprising, since most cacao pathogens did not co-evolve with cacao and are therefore unlikely to have been strong drivers of copy number variation (Bailey and Meinhardt 2018). Exceptions include witches’ broom and, to some extent, frosty pod rot, but even they were not associated with NLR copy number (Meinhardt et al. 2008; Bailey et al. 2018). It is still possible, however, that copy number variation was driven by co-evolution with one or multiple unknown pathogens and/or mutualists (e.g. Mycorrhizae), whose associations with hosts are in part also mediated by NLR genes.

Most variation in NLR content was localized to Chr5, Chr6, Chr7, and Chr10 (Figure 2A), which together accounted for 45-85% of all NLRs, depending on genotype. Those four chromosomes also have the greatest concentration of pseudogenes (Figure 2B). These results are consistent with previous findings that expansion of NLR clusters occurs via a birth-and-death process leading to the formation of NLR hotspots (Meyers et al. 2003; Mizuno et al. 2020). Developed as a counterpoint to models of concerted evolution, the birth-and-death model was originally conceived as an explanation for patterns of evolution observed in the animal major histocompatibility complex (MHC) and immunoglobulin (Ig) gene families (Ota and Nei 1994; Nei et al. 1997). Members of those gene families were more closely related to orthologous genes in other species than they were genes from the same species, inconsistent with expectations under a model of concerted evolution. MHC and Ig genes instead followed a pattern of repeated gene duplication followed by either retention, sometimes for long evolutionary time (Klein 1987), or pseudogenization.

Birth-and-death evolution has now been used to explain the expansion or contraction of many gene families, including the ubiquitin family (Nei et al. 2000), the fatty acyl-CoA reductase family (Finet et al. 2019), and the ATP-binding cassette family (Annilo et al. 2006). Clusters of NLR genes were first shown to evolve through a birth-and-death like process in lettuce (Meyers et al. 1998), but has been confirmed in numerous other species (Jeong et al. 2001; Meyers et al. 2003; Stam, Nosenko, et al. 2019; Mizuno et al. 2020). This same model of birth-and-death evolution was present across our cacao genomes, evidenced by High CNV genotypes possessing a much higher number of NLR pseudogenes than Low CNV genotypes, and a pattern of pseudogene density that exactly mirrored NLR density (Figure 2A-B).

The singular exception to the birth-and-death model was ICS-1, which possessed 2x the number of pseudogenes as it did NLR genes, and 3-4x the number of pseudogenes as the High CNV genotypes (Figure 2B). ICS-1 pseudogenes were localized to the four NLR-dense chromosomes, similar to the High CNV genotypes, but patterns of pseudogenization were different between the two sets. Most pseudogenes in ICS-1 originate from a narrow set of parents, rather than the 1:1 or 2:1 pseudogene:parent NLR relationship in the High CNV genotypes. ICS-1’s unique pattern of NLR abundance could be the result of two possible scenarios. First, ICS-1 may have undergone rapid expansion of NLRs in a manner similar to the high CNV genotypes, but at some point went through a large scale pseudogenization event, likely mediated by retrotransposition (Esnault et al. 2000; Ding et al. 2006). This option, however, seems unlikely given TE abundance was similar across all genotypes (Figure 5 and 6). The second option is that pseudogene expansion may have occurred through the same mechanisms that produced High and Low CNV genotypes, but rather than duplicating functional NLRs, pseudogenes were duplicated (Mighell et al. 2000). This scenario is supported by the pattern of pseudogene parentage outlined above. That is, repeated duplication of a pseudogene would result in a skewed pseudogene:parent NLR ratio, as we observe in ICS-1 relative to the High CNV genotypes. A combination of these two scenarios is also possible, e.g. pseudogenization via retrotransposition followed by pseudogene duplication by unequal crossing over. Ascertaining the respective likelihoods of these scenarios, however, requires further comparative analyses. For instance, if ICS-1 underwent a large-scale pseudogenization event, most of its NLR pseudogenes should be syntenic to functional NLRs in the High CNV genotypes. Likewise, processed pseudogenes formed through retrotransposition should lack introns, assuming the parent genes were not intronless, and have a 3’ poly-A tail (Ding et al. 2006). If ICS-1 pseudogene expansion occurred through duplication of existing pseudogenes, however, intron-exon architecture should be more similar across duplicates than they are to their respective parent NLRs. Interestingly, ICS-1 is susceptible to several cacao pathogens (Figure 3) (Phillips-Mora et al. 2005; Fister et al. 2020) but is also known for its high yield (Araújo et al. 2009). It is tempting to speculate that this difference between defense and yield represents a tradeoff resulting from a combination of demographic history and life history strategy, but further work is required to test this hypothesis.

Differences in NLR copy number appeared to be caused by tandem and proximal duplications that resulted in the formation of NLR clusters. High CNV genotypes had significantly more and larger NLR clusters than Low CNV genotypes (Figure 9B), nearly all of which were localized to the same four chromosomes mentioned above (Figure 9A). This rapid expansion of NLRs could have been caused by two mechanisms. The first is TE-mediated gene duplication, which has a demonstrated role in NLR expansion (Kim et al. 2017). Gene duplication by TEs is most often accomplished by class I transposable elements like LTR and LINE elements (Xiao et al. 2008; Zhu et al. 2016). However, rolling circle Helitrons and DNA Pack-MULEs are also capable of gene duplication (Lai et al. 2005; Jiang et al. 2011). To this end, we investigated the abundance and density of five TE classes across 10 of our genomes. We found no association between TE abundance and NLR abundance (Figure 5 and 6), both when viewing the data in aggregate and when separating TEs by chromosome. Thus, it appears cacao’s rapid expansion of NLRs was not TE-mediated. The other mechanism most likely to generate tandem and proximal duplications is unequal crossing over (Michelmore and Meyers 1998; Leister 2004), which occurs when homologous sequences are incorrectly paired during meiosis. Once tandem and proximal duplicates are formed, the risk of further homologous mismatches increases, resulting in elongation of duplicate arrays and the subsequent expansion of gene families. Indeed, many well-known gene clusters are the result of unequal crossing over, including the human CYP2D6 cluster (Heim and Meyer 1992), the fruit fly glutamate tRNA cluster (Hosbach et al. 1980), and the flax M locus (Anderson et al. 1997). While we did not explicitly test whether unequal crossing over caused the observed patterns of NLR expansion, our results are consistent with this mechanism.

NLR genes are one of the first layers of pathogen defense in plants. Investigating NLR diversity across multiple populations of a single species is therefore necessary to understand how organisms interact with the environment, shaping their ecology, and for harnessing their diversity to breed more resilient crops. However, the biological and technical challenges associated with studying their evolutionary history have resulted in very few investigations into intraspecific NLR variation. Here, we examined the evolution of NLR genes across 11 genotypes of *Theobroma cacao*. Our results indicate local duplications can radically reshape gene families over short evolutionary time scales, creating a source of NLR diversity that could be utilized to enrich our understanding of both plant-pathogen interactions and resistance breeding.

## Data and Resource Availability

Assembled genomes and their corresponding annotations can be found on the Guiltinan-Maximova cacao data repository at the following link: http://bigdata.bx.psu.edu/Cacao_NSF_data/assemblies/Genomes/final_assemblies/10x_meta_assemblies_v1.0. Raw data can be found at the National Center for Biotechnology Information’s Sequence Read

Archive under BioProject PRJNA558793, and can be found at http://identifiers.org/bioproject:PRJNA558793. Custom code used for analyses can be found here: https://github.com/npwinters/Cacao_NLR_Evolution.git

## Acknowledgments

Thanks to Lena Sheaffer for her assistance in project and laboratory management, Paula Ralph for her work extracting genomic DNA. This work was supported by the National Science Foundation Plant Genome Research Program grant IOS-1546863, the US Department of Agriculture National Institute of Food and Agriculture, Federal Appropriations under Project PEN04569 and accession number 1003147, the United States Department of Agriculture National Institute of Food and Agriculture graduate research fellowship (grant no. 2018-07789), The Pennsylvania State University College of Agricultural Sciences, the Huck Institutes of the Life Sciences, and the Penn State Endowed Program in Molecular Biology of Cacao.

## Notes

### Competing Interest Statement

The authors have declared no competing interest.

