## Supplemental Figure 1 for "Local Gene Duplications Drive Extensive NLR Copy Number Variation Across Multiple Genotypes of *Theobroma cacao*"

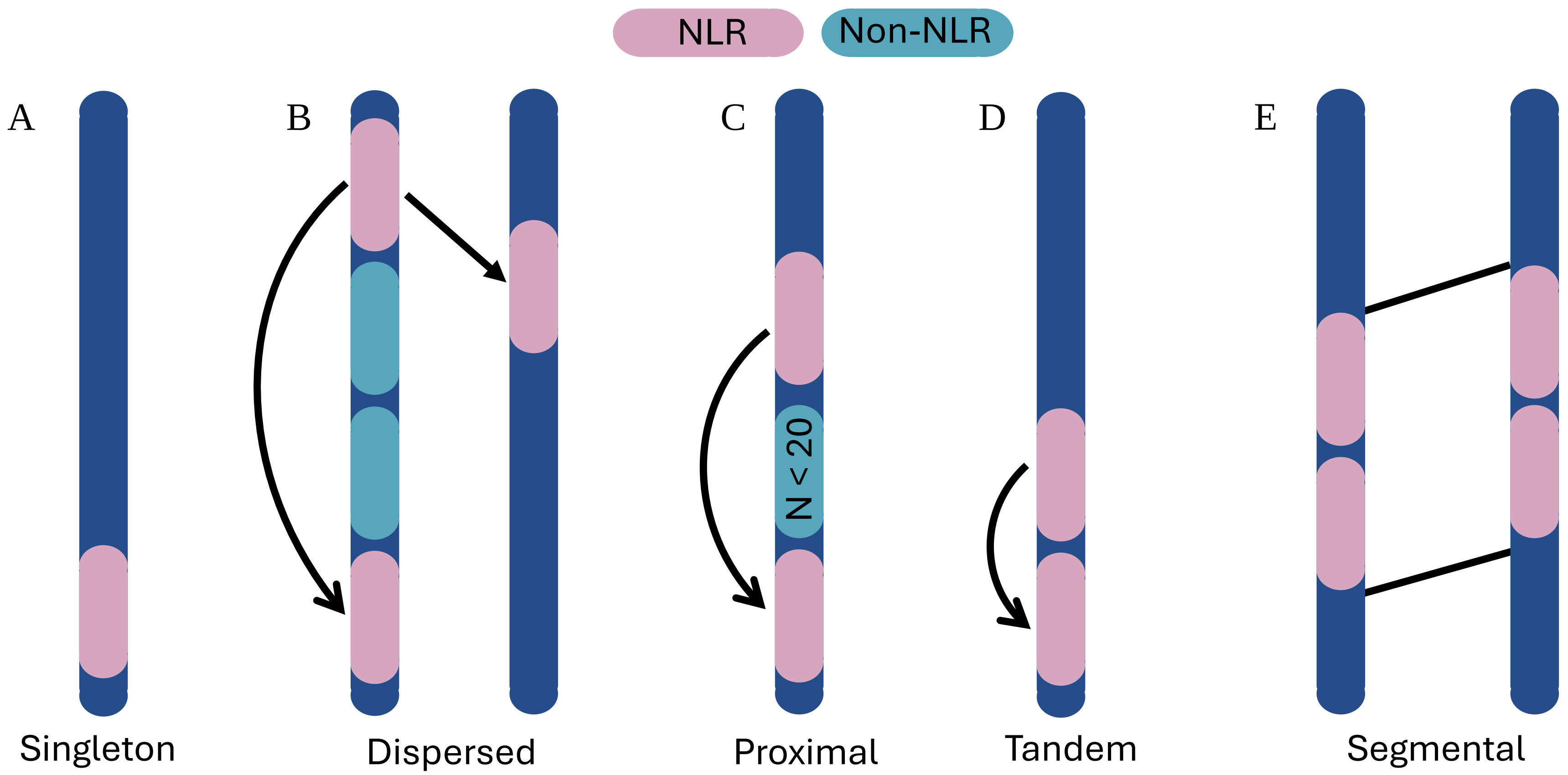


**Supplemental Figure S1: Types of gene duplication.** Duplicate genes were classified into one of five categories: singletons, dispersed, proximal, tandem, or WGD/segmental duplicates. Arrows represent evidence of sequence similarity (BLAST) indicative of a gene duplication. All NLRs were first classified as singletons, i.e. genes with no history of recent duplication (A). If NLR genes contained significant BLAST hits elsewhere in the genome, they were reclassified as dispersed duplicates (B) . Dispersed duplicates were then further categorized as proximal or tandem based on distance between hits. If < 20 genes separated the NLR duplicates, they were considered proximal (C). If NLR duplicates were immediately adjacent to one another, they were considered tandem (D). Lastly, NLR duplicates that were anchors of collinear blocks, as defined by MCScanX's algorithm, were classified as WGD/segmental duplicates (E).
