## Supplemental Figure 2 for "Local Gene Duplications Drive Extensive NLR Copy Number Variation Across Multiple Genotypes of *Theobroma cacao*"

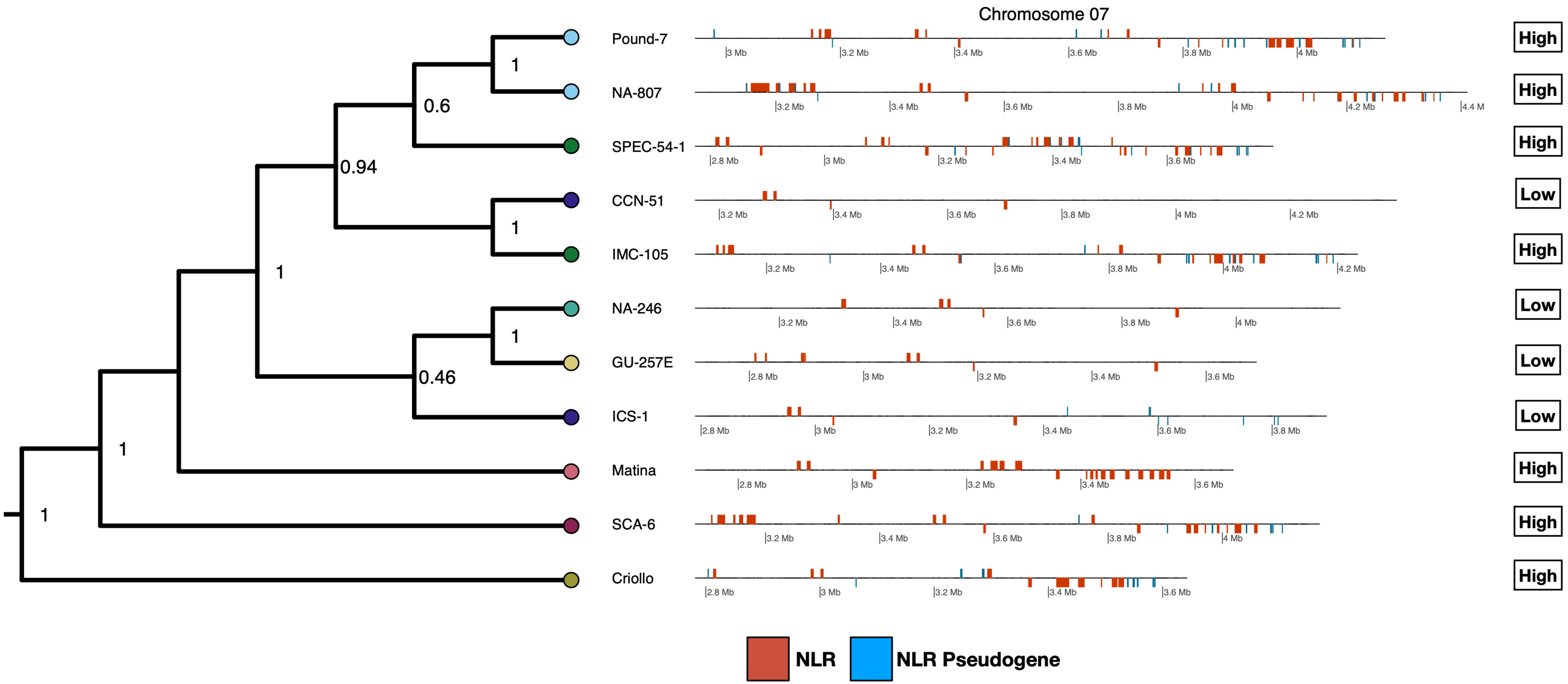


**Figure 10. Synteny of an NLR cluster expanded through local duplications.** NLR genes and NLR pseudogenes are shown as orange and blue bars, respectively. The phylogeny on the left indicates evolutionary relationships between the 11 genotypes used for this study. Labels on the right side indicate whether a genotype is in the Low or High CNV group.
